# Trajectory Mapping of the Early *Drosophila* Germline Reveals Controls of Zygotic Activation and Sex Differentiation

**DOI:** 10.1101/2020.09.11.292573

**Authors:** Hsing-Chun Chen, Yi-Ru Li, Hsiao Wen Lai, Hsiao Han Huang, Sebastian D. Fugmann, Shu Yuan Yang

## Abstract

Germ cells in *D. melanogaster* are specified maternally shortly after fertilization and are transcriptionally quiescent until their zygotic genome is activated to sustain further development. To understand the molecular basis of this process, we analyzed the progressing transcriptomes of early male and female germ cells at the single-cell level between germline specification and coalescence with somatic gonadal cells. Our data comprehensively covered zygotic activation in the germline genome, and analyses on genes that exhibit germline-restricted expression revealed that polymerase pausing and differential RNA stability are important mechanisms that establish gene expression differences between the germline and soma. In addition, we observed an immediate bifurcation between the male and female germ cells as zygotic transcription begins. The main difference between the two sexes is an elevation in X chromosome expression in females relative to males signifying incomplete dosage compensation with a few select genes exhibiting even higher expression increases. These indicate that the male program is the default mode in the germline that is driven to female development with a second X chromosome.

## Introduction

The germline-soma dichotomy is established as germ cells are specified and is central to the unique ability of germline to transition to the totipotent state in embryos (Johnson and Alberio, 2015). In mice, WNT and BMP signaling induce germline formation via the expression of multiple transcriptional regulators such as PRDM14, PRDM1, and TFAP2C (Sybirna et al. 2019). In *D. melanogaster*, germ plasm loaded in the posterior of oocytes contains RNAs and proteins that are important for germline specification and early germline characteristics such as repression of transcription and migration (Santos and Lehmann, 2004). The phase of transcriptional quiescence is thought to prevent erratic expression of somatic genes (Hanyu-Nakamura et al. 2008; Asaoka et al. 2019). However, we do not have a clear understanding of what occurs downstream of specification that embodies the germline program which actively drives the specific development of this lineage. Unlike in mammals, there are no known transcriptional regulators that dictate germline development in the fruit fly.

One important developmental choice germ cells must make early on is to determine their sex. For fruit flies, germline sex requires matching somatic input and intrinsic decision, both of which are set by their sex chromosome compositions (Murray et al. 2010). There are genes known to function intrinsically in regulating germline sex (Hashiyama et al. 2011; Yang et al. 2012), but it is unclear how and when the germline sex is read autonomously. Sex-specific germ cell differentiation is first observed in late embryogenesis, and these events include the division and stratification of male germ cells whereas the female ones remain mostly quiescent (LeBras and Van Doren, 2006; Wawersik et al., 2005). However, gene expression differences that underlie these cellular changes likely take place earlier. Another issue important to sexual differentiation is dosage compensation, a process that occurs to equalize expression of X chromosome genes between the two sexes as females have two doses compared to just one in males. In flies, the mechanism for dosage compensation in somatic cells is well established (Kuroda et al. 2016), but it is not clear if germ cells also require and perform dosage compensation.

The emergence of single-cell RNA-sequencing techniques has opened a new window to investigating critical early germline developmental events especially since germ cells exist in small numbers at these stages. Multiple studies have analyzed transcriptomes of germ cells at the single-cell level, but mostly at later stages of development (Slaidina et al. 2020; Jevitt et al. 2020; Mahadevaraju et al. 2020; Rust et al. 2019; Li et al. 2017; Niu and Spradling 2020). Here we profile the transcriptomes of fruit fly germ cells after specification and before they have coalesced with somatic cells, and we perform this survey for both male and female germ cells. We identify germline-specific genes expressed in this time window, and analyses on the regulatory regions of those activated zygotically reveal that germline-specific expression is likely achieved post-transcriptionally. Moreover, we find that as zygotic transcription starts, a prompt increase in X chromosome expression is observed in females compared to males due to incomplete dosage compensation. Furthermore, a few select genes exhibit higher female-to-male expression ratios, and we provide functional evidence for one of the top candidates, *female sterile (1) homeotic (fs(1)h)*, to be physiologically important for female but not male germline development. Taken together, these indicate that sexual differentiation has begun at this early stage in the germline.

## Results

### Purification of early germ cells for single-cell transcriptome profiling

To obtain germ cells after specification and before gonad coalescence for scRNA-seq, we FACS-purified germ cells from 0-8 h embryos based on germline-specific GFP expression using the *vas-GFP* transgene (Fig. 1a-b, Fig. S1a-d). We also performed a second round of scRNA-seq for which we processed 5-8 h female and male germ cells separately to achieve greater resolution between the sexes (Fig. S1f-i). This was made possible by the addition of a *Sxl-GFP* transgene which is highly expressed in female but not male 5-8h embryos (Fig. S1e). These two female and male samples contained some somatic cells due to GFP expression from the *Sxl-GFP* transgene, but this did not pose a problem for bioinformatics analyses as there are well-established markers that can help us unequivocally identify germ cell populations.

**Figure S1.**
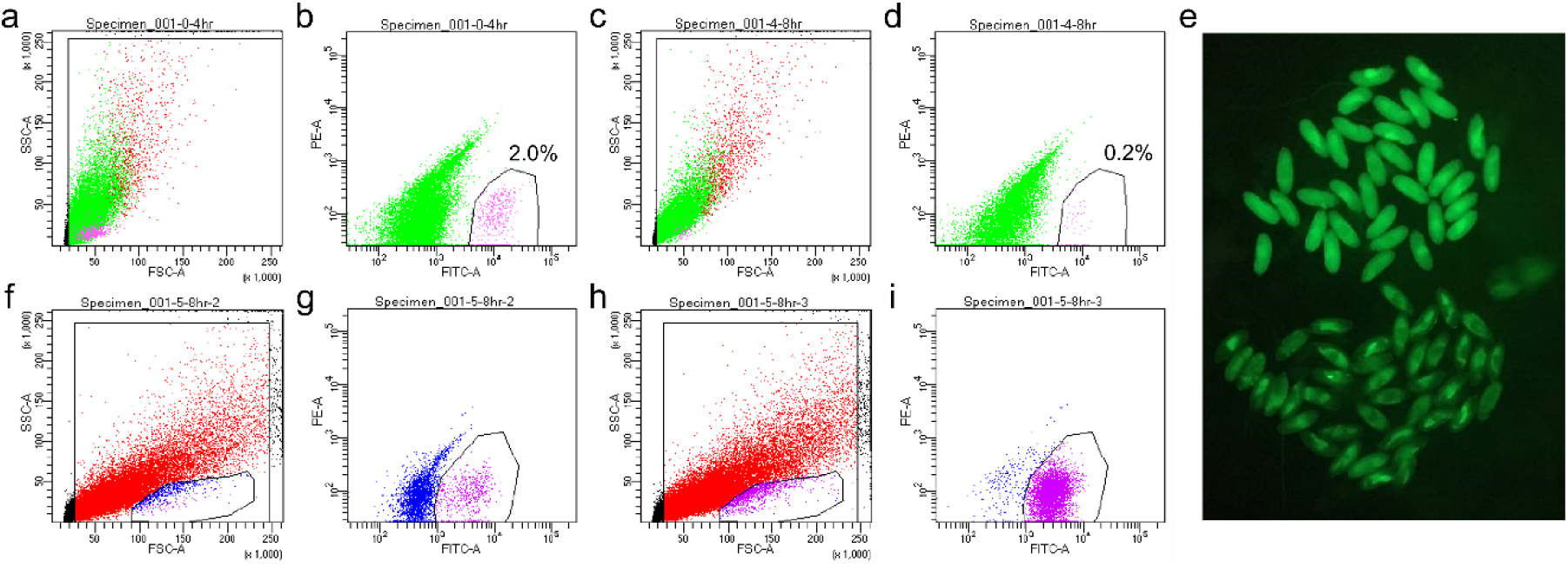
FACS schemes for the collection of single germ cells for scRNA-seq. **a-b**, FACS plots for unsexed 0-4 h germ cells. **c-d**, FACS plots for unsexed 4-8 germ cells. **a, c**, Gating for live cells which plots FSC-SSC. **b, d**, The gate for the magenta cells indicate the GFP+ cells that were sorted and their percentages. X and Y axes indicate the FITC and PE channels, respectively. **e**, Female and male 5-8 h embryos can be separately based on their *Sxl-GFP* expression under a fluorescent stereoscope. The embryos in the upper half express GFP highly and are females; those in the lower half are males. **f-g**, FACS plots for female 5-8 h germ cells. **h-i**, FACS plots for male 5-8 h germ cells. **f, h**, The small encircled populations on the bottom of the plots are enriched for germ cells, thus this live gate was used for the FITC-PE plots (**g, i**). **g, i**, FITC-PE plots with magenta cells being the GFP+ cells sorted. The axes are the same as in **b, d**.

**Figure 1.**
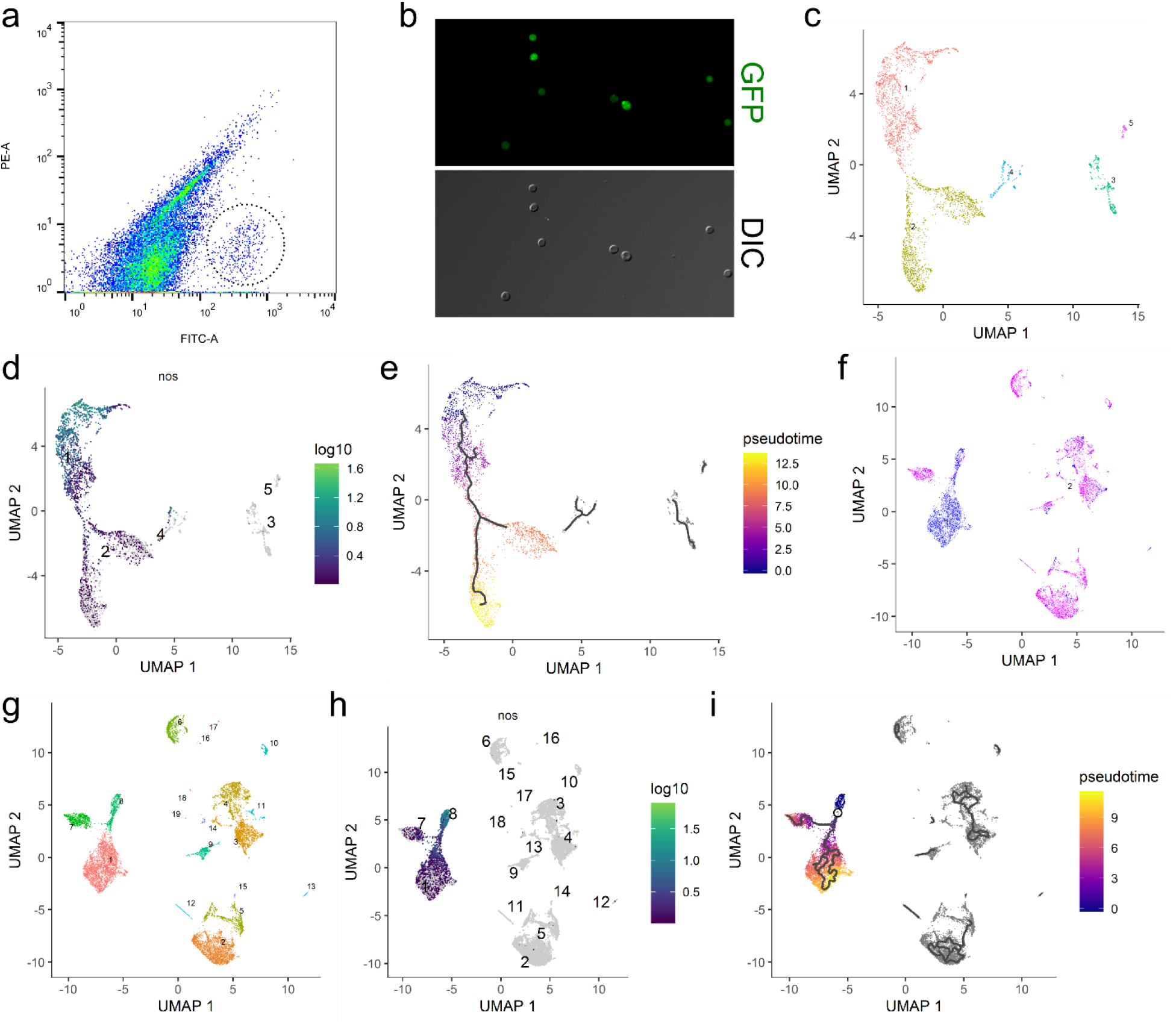
Clustering and pseudotime analysis of single-cell germline transcriptomes. **a**, FACS plot for sorting germ cells from embryonic homogenates of 0-8 h embryos with the GFP+ germ cells highlighted by a dotted circle. X-axis is the green channel. **b**, Morphology by DIC and GFP profile of sorted 0-8 h germ cells in the unsexed sample. Note that virtually all cells recovered are GFP-positive cells whose morphology resemble that of germline. **c**, Clustering analysis of the unsexed sample. Each dot represents a cell in the dataset, and individual clusters are numbered and colored differently. **d**, Expression profile of nos of the unsexed sample showing that clusters 1 and 2 make up the main germline cluster. Each dot represents a cell in the dataset, the numbers indicate individual clusters, and the color code for expression levels is indicated on the right. **e**, Pseudotime analysis of the germline cluster from the unsexed sample. The black lines within the clusters indicate pseudotime trajectories. The legend on the right explains the color code of pseudotime. **f**, The sexed dataset is plotted after dimension reduction. Each blue and magenta dot represents a cell from the male and female dataset, respectively. **g**, Clustering analysis of the sexed sample. Each dot represents a cell in the dataset, and individual clusters are numbered and colored differently. **h**, Expression profile of *nos* after clustering analysis of the sexed dataset indicating that clusters 1, 7, and 8 contain germ cells. Details of the graph is the same as in **b. i**, Pseudotime analysis of the germline cluster from the sexed dataset. Black lines indicate pseudotime trajectories and the color codes of pseudotime is indicated on the right.

### Clustering and pseudotime analysis reveal bifurcation between early male and female germline

The primary bioinformatics tool we used for scRNA-seq data analysis was Monocle3 which enables both clustering and pseudotime analyses (Cao et al. 2019). Clustering analysis of the sequencing data from 0-8 h unsexed cells resulted in a major Y-shaped germline cluster based on the expression profiles of known germline marker genes such as *nanos (nos)* and *vasa (vas)* (Fig. 1c-d, Fig. S2a-b). The female and male samples, when combined (herein referred to as the “sexed sample/dataset”), also produced a Y-shaped germline cluster as evidenced by the profiles of *nos* and *vas* expression (Fig. 1g-h, Fig. S2e-f).

**Figure S2.**
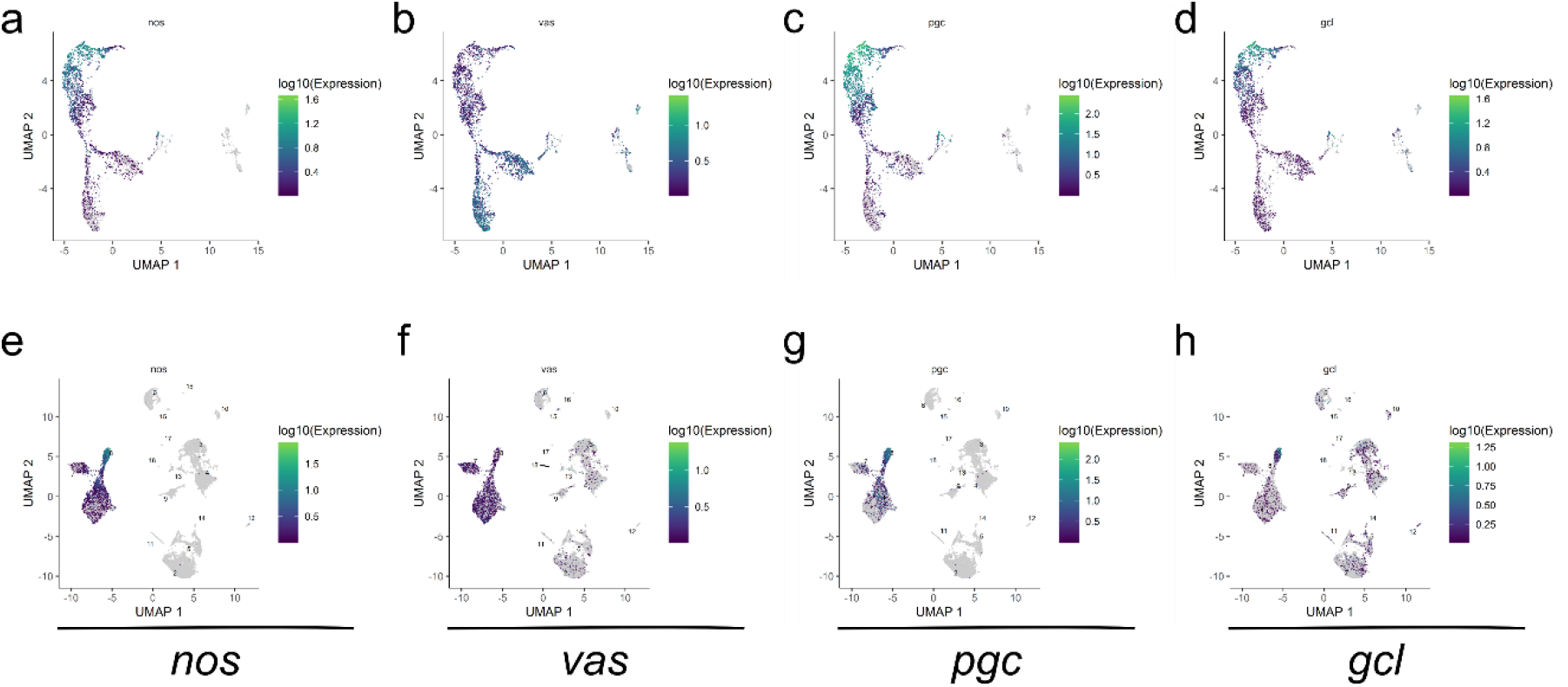
Expression profiles of known germline markers. **a-d**, Unsexed sample. **e-h**, Sexed sample. **a**, **e**, *nos* expression. **b**, **f**, *vas* expression. **c**, **g**, *pgc* expression. **d**, **h**, *gcl* expression. Color codes for expression levels are to the right of each plot.

The Y-shaped nature of the germline clusters suggests that the early germ cells we collected are on a transcriptional continuum that diverge once during this time period. When we examined known maternally-contributed germline genes such as *polar granule component* (*pgc*) and *germ cell-less* (*gcl*) (Nakamura et al. 1996; Jongens et al. 1992), we found that their transcript levels are highest in cells at the end of the Y-stems and gradually decrease towards the branches of the germline clusters (Fig. S2c-d, g-h). This indicates that the earliest germ cells are at the base of the Y-stems, a position assigned as the “root” of the clusters for pseudotime analyses, which subsequently gave rise to trajectories along pseudotime which start at the base of Y-shape, branch in the middle, and progress towards the tips of the two prongs (Fig. 1e, i). In the sexed sample, the stem contains germline from both sexes whereas the branches are comprised largely of either male or female germline (Fig. 1f, i). This indicates that sexual differentiation has begun in this early time point prior to gonad coalescence.

### Distinct germline expression patterns along pseudotime

The presence of distinct expression patterns common to many genes are suggestive of developmental changes and transitions between physiological states. Of the multiple expression modules determined for the germline clusters, several exhibited a clear and continuous trend over pseudotime and are shared between the unsexed and sexed samples (Fig. S3a-b).

**Figure S3.**
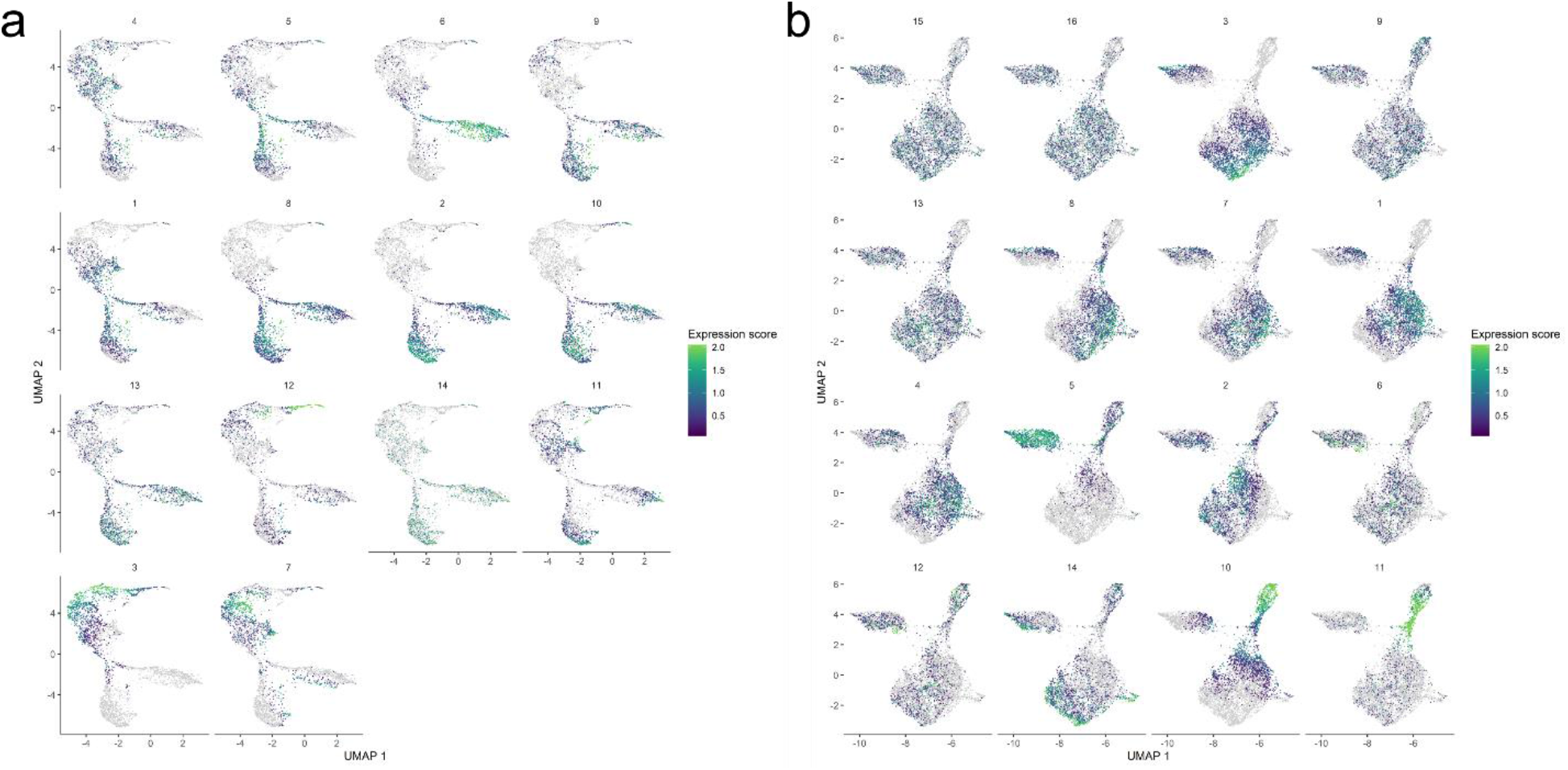
Expression modules for the germline clusters in the unsexed (a) and sexed (b) datasets. The module numbers are indicated on top of each individual module, and color codes for expression levels are indicated on the right.

One category, exemplified by module 3 of the unsexed sample and module 10 of the sexed sample, represents genes that are mainly expressed maternally (Fig. 2a, d). Another type portrays increased gene expression along pseudotime, reflecting zygotic activation of the germline genome, and includes module 2 from the unsexed sample and module 3 from the sexed sample (Fig. 2b, e).

**Figure 2.**
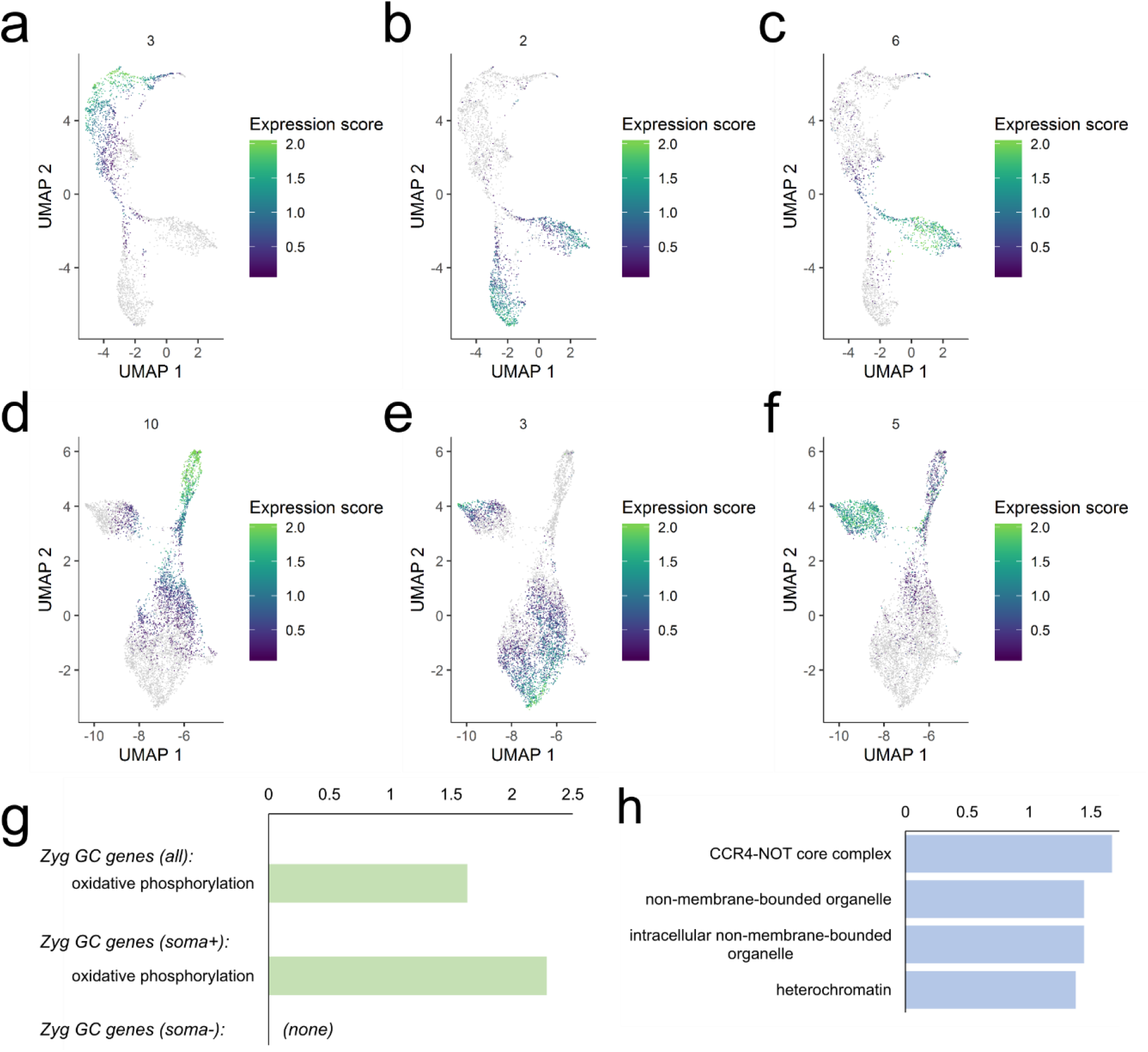
Expression trends in the germline clusters and activation of the zygotic germline genome. **a-f**, Expression modules in the unsexed (**a-c**) and sexed (**d-f**) samples indicating patterns of maternal contribution (**a, d**), zygotic activation (**b, e**), and female-biased expression (**c-f**). The color codes for expression levels are to the right of each graph. **g**, KEGG pathway enrichment results for the genes activated in the zygotic germline. The upper bar shows the enrichment term found for all zygotic genes together while the lower bar is for those that are soma-positive. There were no terms found to be enriched for the soma-negative genes. **h**, GO-Term: Cellular Components enrichment analysis for the germline-specific zygotic genes.

Notably, there are also modules that show higher expression in one branch than the other (module 6 from unsexed sample and module 5 from the sexed sample, Fig. 2c, f), and we know from the sexed sample that the branch with higher expression is comprised of female germ cells (Fig. 1f). Intriguingly, there are no modules with the reverse trend.

We next determined “marker genes” for the germline population from the unsexed sample before and after the bifurcation point to find genes that are maternally-deposited versus zygotically-produced (Table S1, “Maternal” and “Zygotic”). We also picked out a subset of markers that are highly germline-enriched by removing those that showed substantial somatic expression as determined by the somatic cells included in the sexed sample (Table S1, “Maternal soma-negative”, “Zygotic soma-negative”).

The gene lists were subsequently validated with the embryonic *in situ* hybridization database of the Berkeley Drosophila Genomic Project (BDGP, insitu.fruitfly.org/cgi-bin/ex/insitu.pl). We examined the top 25 candidates for both the “Maternal soma-negative” and the “Zygotic soma-negative” groups of genes. We find that for both lists the *in situ* patterns of most genes which have been profiled at BDGP match the expectations based on our sequencing results (Fig. S4a). Of the ones that do not match, most gave no *in situ* signals which may be a sensitivity issue. We compiled example images of one gene each from the maternal and zygotic lists to demonstrate the similarities between the BDGP results and ours (Fig. S4b-g). We also examined the *in situ* patterns of two zygotically activated candidates not profiled by BDGP, one belonging to the soma-negative group (*HP6*) and the other in the soma-positive group (*P32*). We find that their expression indeed gradually increases in the embryonic germline (Fig. S4h-l). Overall there is very good accordance between our two rounds of scRNA-seq and the BDGP *in situ* patterns, indicating that our transcriptome results are robust.

**Table S1.** Maternal and zygotic germline markers. (Truncated version; see complete version online.)

| Maternal all | Maternal soma- | Zygotic all | Zygotic soma- |
| --- | --- | --- | --- |
| pgc | pgc | CG17650 | CG17650 |
| wisp | wisp | CG32971 | CG32971 |
| dhd | dhd | CG14346 | CG14346 |
| nos | nos | CG43088 | CG43088 |
| 26-29-p | 26-29-p | CG18605 | CG18605 |
| alphaTub67C | alphaTub67C | ATPsynepsilonL | ATPsynepsilonL |
| CanB2 | CanB2 | HP6 | HP6 |
| Jafrac1 | mtrm | CG12477 | CG12477 |
| Stlk | rpk | RpS10a | RpS10a |
| mtrm | BigH1 | CG42857 | CG42857 |
| rpk | CG8507 | CG4415 | CG4415 |
| BigH1 | ldh | qjt | qjt |
| Mdh2 | PyK | CG14932 | CG14932 |
| CG8507 | CG1927 | P32 | gnu |
| CycB | shu | gnu | hog |
| Echs1 | CG6287 | CG4004 | CG32706 |
| ldh | CG17734 | hog | CG34261 |
| MED26 | plu | CG32706 | Oxp |
| Atf6 | Pdk | CG34261 | Rcd-1r |
| PyK | Drep1 | IscU | RpL22-like |
| CG1927 | Pxt | Oxp | Rpt3R |
| retn | GstT1 | Rcd-1r | CG15262 |
| shu | Cen | Fkbp59 | CG5194 |
| CG6287 | Gapdh1 | RpL22-like | CG14864 |
| CG17734 | Cyp6v1 | GLaz | CG14757 |
| plu | Npc2a | Rpt3R | blanks |
| Pdk | Z600 | CG3176 | Cyt-c1L |
| CG12112 | Gip | CG15262 | CG6999 |
| Sesn | CG11674 | CG5194 | agt |
| Drep1 | Jabba | lncRNA:CR43356 | CG14036 |
| Tao | Uch | Chmp1 | CG15398 |
| Eno | BicD | CG32625 | karr |
| me31B | Gabat | CG5039 | Dhfr |
| CG7115 | ebd2 | CG14864 | CG1894 |
| Pxt | yip2 | CG32486 | Sdhaf3 |
| GstT1 | del | CG1890 | CG11125 |
| dnc | AGO3 | CG15536 | Tspo |
| Sod3 | CG4404 | CG7878 | CG17625 |
| Pino | CG4586 | mRpS34 | HP1D3csd |

### Activation of the zygotic germline genome

Of the genes activated zygotically in the germline, some are highly germline-enriched but the majority are also expressed in the soma, albeit at lower levels. When we looked at the molecular functions of these two types of genes, those that are also expressed somatically are mostly “house-keeping” genes while those that are more germline-specific are related to germline-enriched features such as RNA granules (Fig. 2g-h, Table S2). Interestingly, zygotic germline genes are most enriched for mitochondrial components (Fig. 2g), an observation also made for the embryonic human germline (Li et al. 2017). This may reflect a conserved requirement for energy consumption in the early germline.

**Figure S4.**
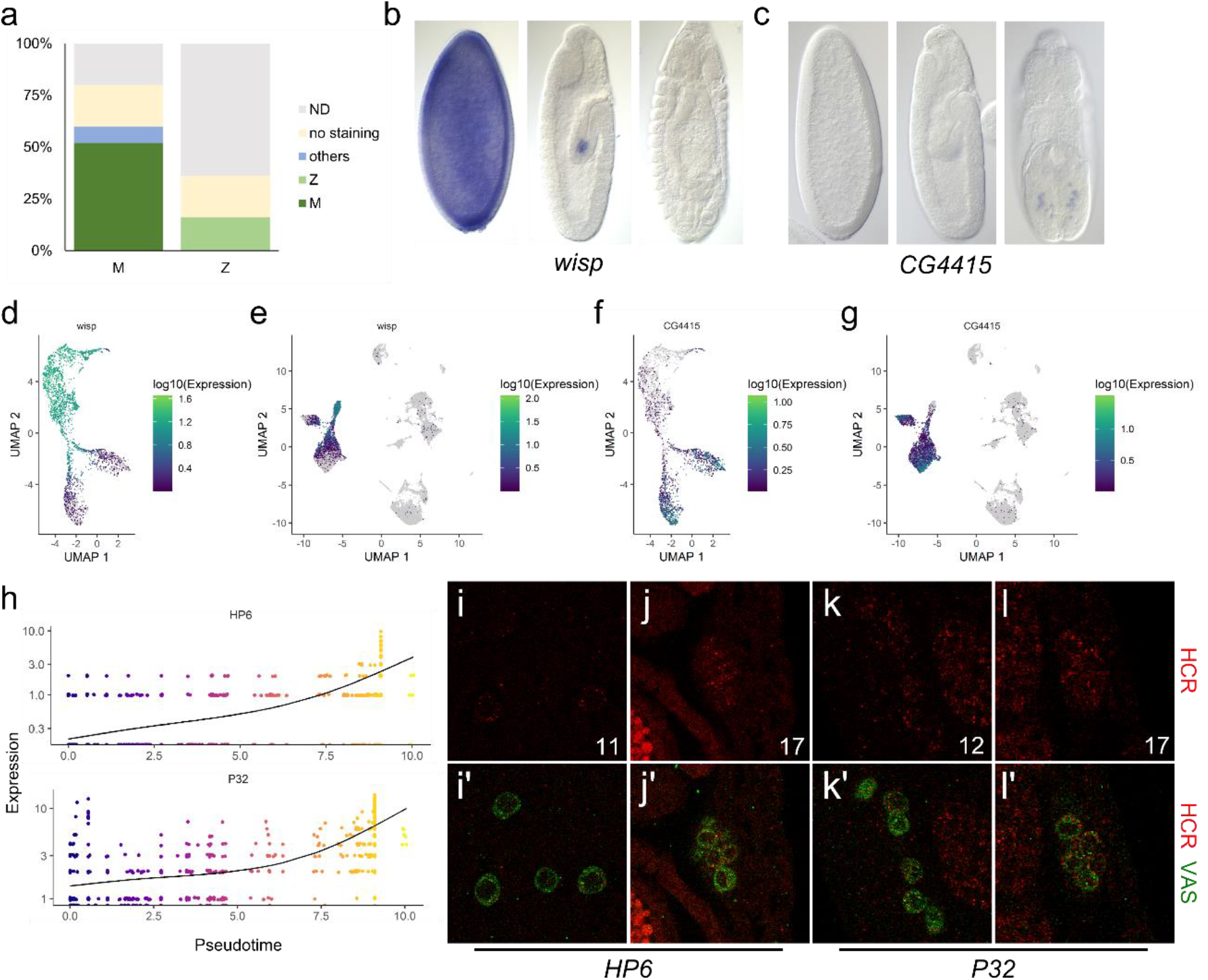
Expression validation of the scRNA-seq results. **a**, Expression patterns of the top 25 germline marker genes in the BDGP database. The left bar is for maternally-deposited, germline-specific genes (M). The bar on the right is for zygotically-activated, germline-specific genes (Z). Dark green indicates highest expression in stages 4-6 whereas light green indicates germline-specific expression that begins in stages 7-10 and increases in later stages. Blue indicates that the BDGP in situ pattern was mainly outside of germ cells. Yellow indicates no staining. Gray indicates the fractions that have not been profiled in the database. **b-c**, In situ images from the BDGP database of progressive embryonic stages from left to right for *wisp* (**b**) and *CG4415* (**c**), from the “maternally-deposited” and “zygotically-activated” categories. C, *In situ* images of progressive embryonic stages from left to right from the BDGP database for *CG4415*, a gene from the “zygotic GC” category. **d-e**, Expression profiles of *wisp* in the unsexed (**d**) and sexed combined (**e**) datasets. **f-g**, Expression profiles for *CG4415* in the unsexed (**f**) and sexed combined (**g**) datasets. **h**, Expression levels of *HP6* (upper graph) and *P32* (lower graph) of every cell in the common stem and male branch in the germline cluster of the unsexed sample is graphed along pseudotime. The black lines indicate the trends of expression along pseudotime. The X- and Y-axes plot pseudotime and expression levels, respectively, and the pseudotime color codes are indicated to the right. **i-l**, Fluorescent in situ HCR images for *HP6* (**i-j**) and *P32* (**k-l**) showing expression at stage 17 (**j, l**) and the stages we could detect the first signs of expression (stage 11 for *HP6*, panel **i**, and 12 for *P32*, panel **k**). **i-l** display the HCR channel alone (red) while **i’-l’** display merged images with α-Vasa staining (green) to mark the germ cells.

**Table S2.** GO-term analysis of soma-positive zygotic germline genes. (Truncated version; see complete version online.)

| Source | Term_name | Term_id | Adjusted_p_value |
| --- | --- | --- | --- |
| GO:CC | mitochondrial protein complex | GO:0098798 | 1.16E-47 |
| GO:CC | intracellular membrane-bounded organelle | GO:0043231 | 6.05E-43 |
| GO:CC | protein-containing complex | GO:0032991 | 5.79E-41 |
| GO:CC | membrane-bounded organelle | GO:0043227 | 9.27E-41 |
| GO:CC | intracellular | GO:0005622 | 7.93E-38 |
| GO:CC | mitochondrion | GO:0005739 | 6.76E-35 |
| GO:CC | intracellular organelle | GO:0043229 | 9.76E-35 |
| GO:CC | organelle | GO:0043226 | 4.14E-33 |
| GO:CC | cytoplasm | GO:0005737 | 5.62E-30 |
| GO:CC | membrane-enclosed lumen | GO:0031974 | 9.13E-28 |
| GO:CC | intracellular organelle lumen | GO:0070013 | 9.13E-28 |
| GO:CC | organelle lumen | GO:0043233 | 9.13E-28 |
| GO:CC | mitochondrial ribosome | GO:0005761 | 1.90E-27 |
| GO:CC | organellar ribosome | GO:0000313 | 1.90E-27 |
| GO:CC | respiratory chain complex | GO:0098803 | 1.69E-24 |
| GO:CC | ribonucleoprotein complex | GO:1990904 | 1.70E-24 |
| GO:CC | mitochondrial respirasome | GO:0005746 | 2.81E-23 |
| GO:CC | respirasome | GO:0070469 | 4.18E-23 |
| GO:CC | inner mitochondrial membrane protein complex | GO:0098800 | 8.46E-22 |
| GO:CC | mitochondrial matrix | GO:0005759 | 1.50E-20 |
| GO:CC | mitochondrial respiratory chain complex I | GO:0005747 | 6.84E-18 |
| GO:CC | respiratory chain complex I | GO:0045271 | 6.84E-18 |

We next wanted to investigate whether mechanisms of transactivation are different between soma-negative germline genes, soma-positive germline genes, and genes specific to somatic cell types. To address this point, we looked for enrichment of known transcription factor (TF) binding sites and also performed *de novo* discovery of enriched sequence motifs in the promoter regions of zygotic germline genes (Fig. 3a-b). The distribution of the motifs identified by both strategies is concentrated close to transcription start sites and conforms to what is commonly observed for TF binding sites (Fig. 3c). Intriguingly, their presence on soma-negative and positive germline genes are similar (Fig. 3d), suggesting that these two groups of genes are activated similarly. To compare, we looked for TF binding site enrichment in three somatic clusters in our sexed dataset (clusters 2-4 in Fig. 1g). We found that while the motifs enriched in germline genes are also present in marker genes of somatic clusters (Fig. 3d), their frequencies are reduced and the TF binding sites that are most significantly enriched in the soma are distinct from those in the germ cells (Fig. 3a). Overall, these results do not rule out possible roles for germline-specific TFs in germline-specific expression. Nonetheless, they clearly suggest that transactivation of zygotic germline genes with or without soma expression is quite similar, and the transcription factors likely involved are not restricted to acting in the germ cells.

**Figure 3.**
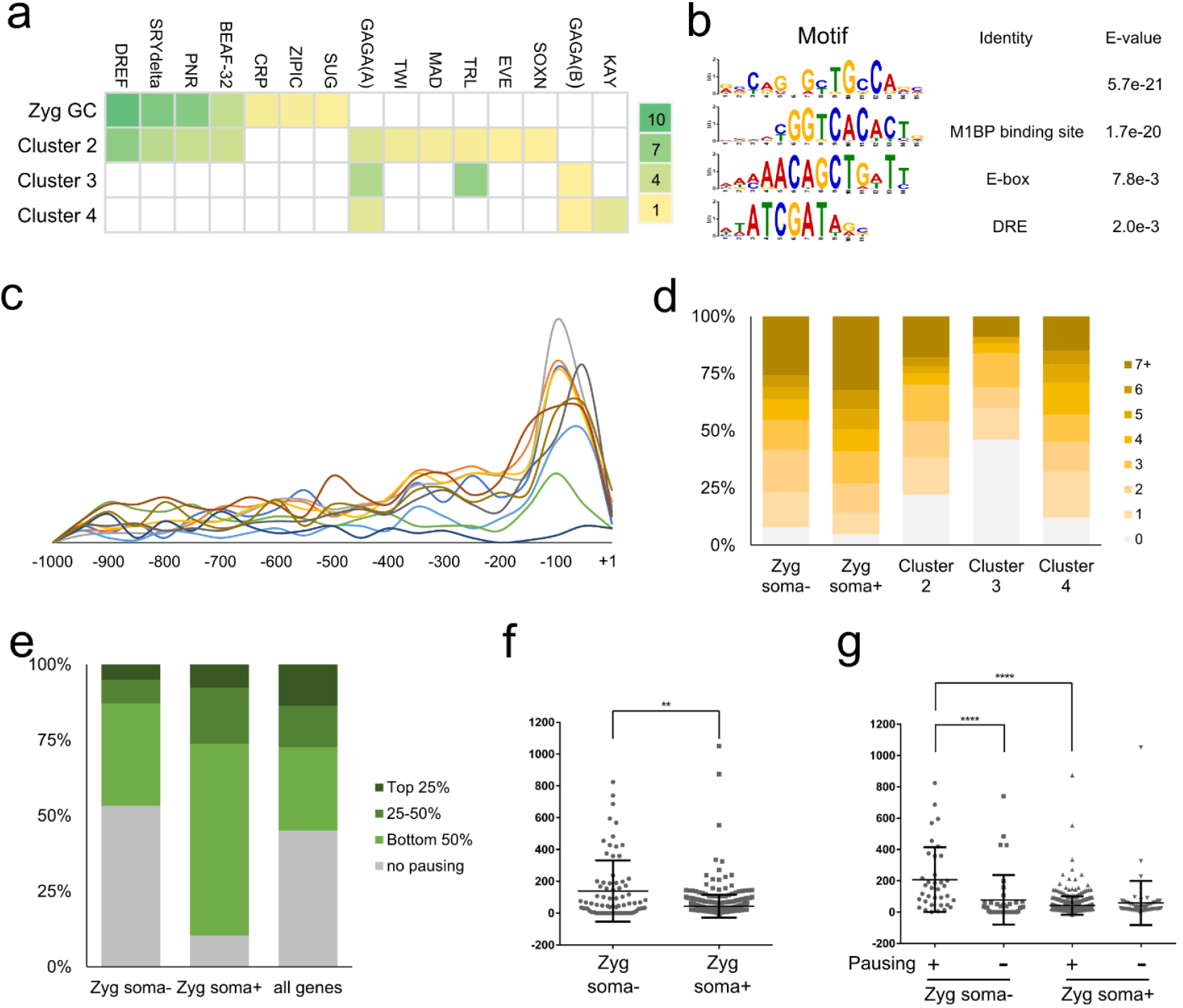
Regulation of early germline specific expression. **a**, Heat map display of transcription factor binding site enrichment analysis for all zygotic germline genes combined and markers of somatic clusters 2-4. The names of TFs are listed on the top and the scale on the right indicates the −log(*p*) values for each entry. **b**, Enriched motifs discovered de novo in the promoter regions of zygotic germline genes and their likely identity based on comparisons with consensus sequences of known TFs. DRE is the binding site of DREF, a factor whose binding site was found to be enriched in (**a**). **c**, Distribution of TF sites enriched in the promoter regions of zygotic germline genes. Each line represents a different enriched motif. The x-axis indicates the position in relations to the transcription start site (+1). **d**, Numbers of binding sites found in the promoter regions of zygotic germline genes or markers of somatic clusters for TFs enriched in the germline. **e**, Polymerase pausing of zygotic germline genes in the early embryonic soma based on a study from Saunders et al. Genes are categorized based on their pausing index into top 25%, 25-50%, bottom 50%, or no pausing groups. We present the comparisons between soma-negative zygotic germline genes, soma-positive zygotic germline genes, and all genes in the genome. **f**, Test of RNA stability in the soma using ratios of nascent RNA levels in the gene body regions to steady-state RNA levels of various genes. The middle horizontal lines indicate the means calculated for the soma-negative and soma-positive groups of genes whereas the top and bottom horizontal lines mark the standard deviations. ** indicates *p* <0.005. **g,** Comparisons of RNA stability between the soma-negative and positive zygotic germline genes that exhibit pausing and not. The values are calculated as in (**f**). **** denotes *p* <0.0001.

### Polymerase pausing and regulated RNA stability contribute to germline-specific expression

If germline-enriched expression is not the result of germline-restricted transcriptional activation, there would need to be additional mechanisms post-transactivation to confer germline-specific gene expression, such as inhibition of transcription elongation and regulation of transcript stability. Here we take advantage of a genome-wide nascent RNA-sequencing (GRO-seq) study in the soma of early *Drosophila* embryos carried out by Saunders et al. to investigate the aforementioned possibilities (Saunders et al. 2013). In the GRO-seq study, pausing of RNA polymerase II after transcriptional initiation was observed for more than half of all genes in somatic cells, and this phenomenon was suggested as a strategy to accommodate greater flexibility in gene expression control during development.

By referencing their results, we find that a subset of zygotic germline genes, both soma-negative and positive, exhibit promoter-proximal polymerase pausing (Fig. 3e). This suggests that transcription of a substantial fraction of zygotic germline genes are indeed activated somatically but experience polymerase pausing which dampens expression of these genes in the soma.

We further used the GRO-seq data to investigate whether differential RNA stability also regulates expression of germline genes in the soma. We determined RNA stability by calculating the ratio of nascent RNA levels in the gene body of various genes from the GRO-seq study to steady-state RNA levels detected in the somatic cells profiled in our dataset. Increases in this ratio would indicate greater RNA instability. Interestingly, we found that RNA turnover in soma-negative zygotic germline genes is greater than that in soma-positive ones; the same trend also holds true when we look specifically at genes that exhibit polymerase pausing (Fig. 3f-g). This suggests that faster turnover rates of transcripts is in part why RNAs of soma-negative genes fail to accumulate despite being transcribed. Interestingly, transcripts of genes that exhibit polymerase-pausing are less stable than those that do not pause (Fig. 3g). These findings indicate that promoter pausing and regulation of RNA stability are important mechanisms that mediate germline-specific gene expression and contribute to establishing the distinct programs between germline and soma.

### Potential germline-enriched factors that regulate zygotic activation in the germ cells

From our scRNA-seq datasets, several maternal and zygotic germline-specific genes known to or potentially encode RNA- and DNA-binding proteins were identified, and they are candidates that can mediate germline-specific expression via transcriptional activation, elongation, RNA stabilization, or translational control (Table S3). Those that are maternally contributed are expressed prior to zygotic activation and are much more likely to be involved in regulating germline-specific zygotic expression. The zygotic ones may contribute to germline-specific gene expression or functions at later stages.

**Table S3.** Early germline specific genes that encode RNA binding proteins or transcription factors.

| **Maternal** |  |
| --- | --- |
| RNA binding proteins: | *nos* |
|  | *osk* |
|  | *bru1* |
| Transcription factors/Zinc fingers proteins: | *cad* |
|  | *ebd2* |
|  | *CG4404* |
|  | *dl* |
|  | *CG9925* |
|  | *tll* |
| **Zygotic** |  |
| RNA binding proteins: | *CG32706* |
|  | *CG6999* |
| Transcription factors/Zinc fingers: | *CG34031* |
|  | *CG14036* |
|  | *CG3919* |
|  | *CG14135* |

### Germline X chromosome expression is partially dosage compensated

Our clustering and pseudotime analyses indicate that as zygotic transcription is activated, the transcriptomes of female and male germ cells diverge (Fig. 1e, i). In addition, there are only expression modules in which female expression is substantially higher than that of males but not the reverse (Fig. S3). Remarkably, we found that genes belonging to these module of female-enrichment are almost all on the X-chromosome and are distributed across the entire X-chromosome (Fig. 4a-b). The observation that expression of many X-chromosome genes are higher in female germ cells than males brings to our mind the issue of dosage compensation. Previous reports have suggested that usage and expression of the X chromosome in the germline could be stage-dependent (Gan et al. 2010; Shi et al. 2020; Mahadevaraju et al. 2020; Hense et al. 2007; Kemkemer et al. 2011; Meiklejohn et al. 2011).

**Figure 4.**
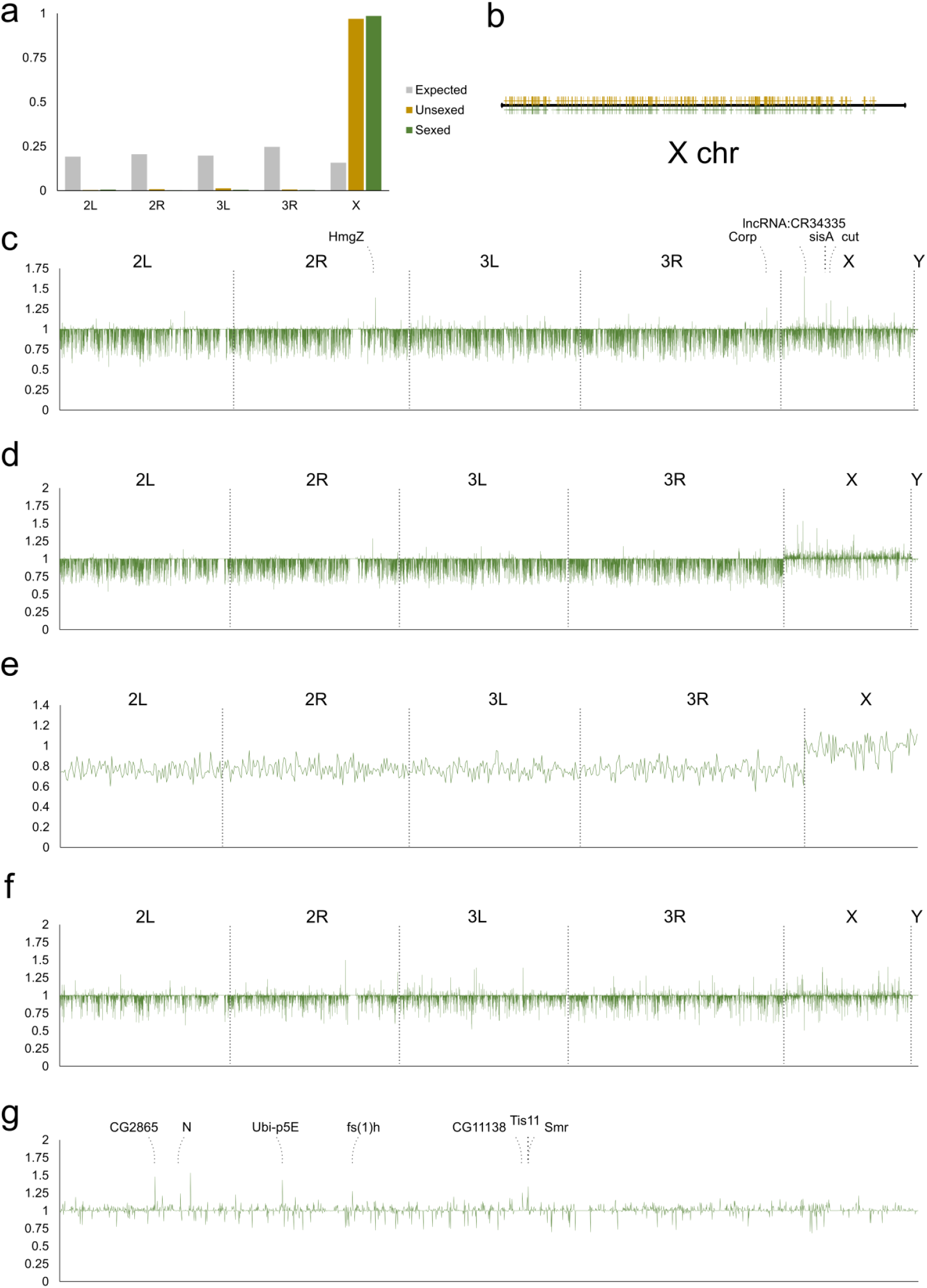
Analyses of sex differences in the early germline. **a**, Chromosomal distribution of genes from the female-enriched expression modules. Grey bars, expected distribution based on number of genes on each chromosome; yellow bars, distribution of genes from the unsexed sample; green bars, distribution of genes from the sexed sample. **b**, Distribution of X-chromosome genes from female-enriched expression modules along the X chromosome. The yellow and green marks designated genes from the unsexed and sexed datasets, respectively. **c**, Differences in expression levels of all genes between female and male germ cells prior to the bifurcation point of the germline cluster from the sexed dataset. The X- and Y-axes are the same for **c-f**: the X-axis plots the positions on individual chromosomes as indicated and the Y-axis plots the fold differences of expression of female germ cells over male germ cells. **d**, Differences in expression levels of all genes between female and male germ cells after they bifurcate in the germline cluster from the sexed dataset. **e**, Expression differences between the two sexes for the 647 zygotically activated germline genes. **f**, Female-to-male expression ratio of all genes of somatic cells included in the sexed sample. **g**, Female-to-male expression ratios of X-chromosome genes of germ cells after bifurcation from the sexed dataset. The identities of the highest peaks are indicated.

To address this point in the embryonic germline, we compared sex differences in autosomal and sex chromosomal expression in the germline before and after the bifurcation point so as to track changes that may occur as a result of zygotic transcription and possible dosage compensation. By plotting the ratio of female-to-male gene expression on different chromosomes, we observed two striking patterns. First, X chromosome gene expression in the females is on average higher than in males, and this phenomenon is specific to the X chromosome (Fig. 4d, Fig. S5a). Second, this sex difference in X chromosome expression occurs only after zygotic transcription begins (Fig. 4c, d). While there is substantial gene-to-gene variation in expression differences, presumably due to the inherently noisy nature of single cell transcriptome data, the overall trend is consistent and the average difference of all X chromosome genes compared to autosomes is about 5%. However, as these genes could be expressed maternally and/or zygotically, it is important to focus on those that are only zygotically activated for a more accurate analysis of the possible effects of dosage compensation. Hence we re-plotted the female-to-male expression ratios for the 647 zygotically activated genes (Table S1) and found that the average expression on the X in females is now about 30% higher than that in males (28% for the sexed dataset, 37% for the unsexed dataset, Fig. 4e, Fig. S5b). To compare, we also looked at the sex expression ratios in somatic cells which are known to be dosage-compensated. This was done using data from the somatic cells collected in the sexed sample, and we found no chromosome-wide differences between expression on autosomes and the X-chromosome (Fig. 4f). Taken together, our results demonstrate that there is partial, incomplete dosage compensation for the X chromosome in the early germline.

**Figure S5.**
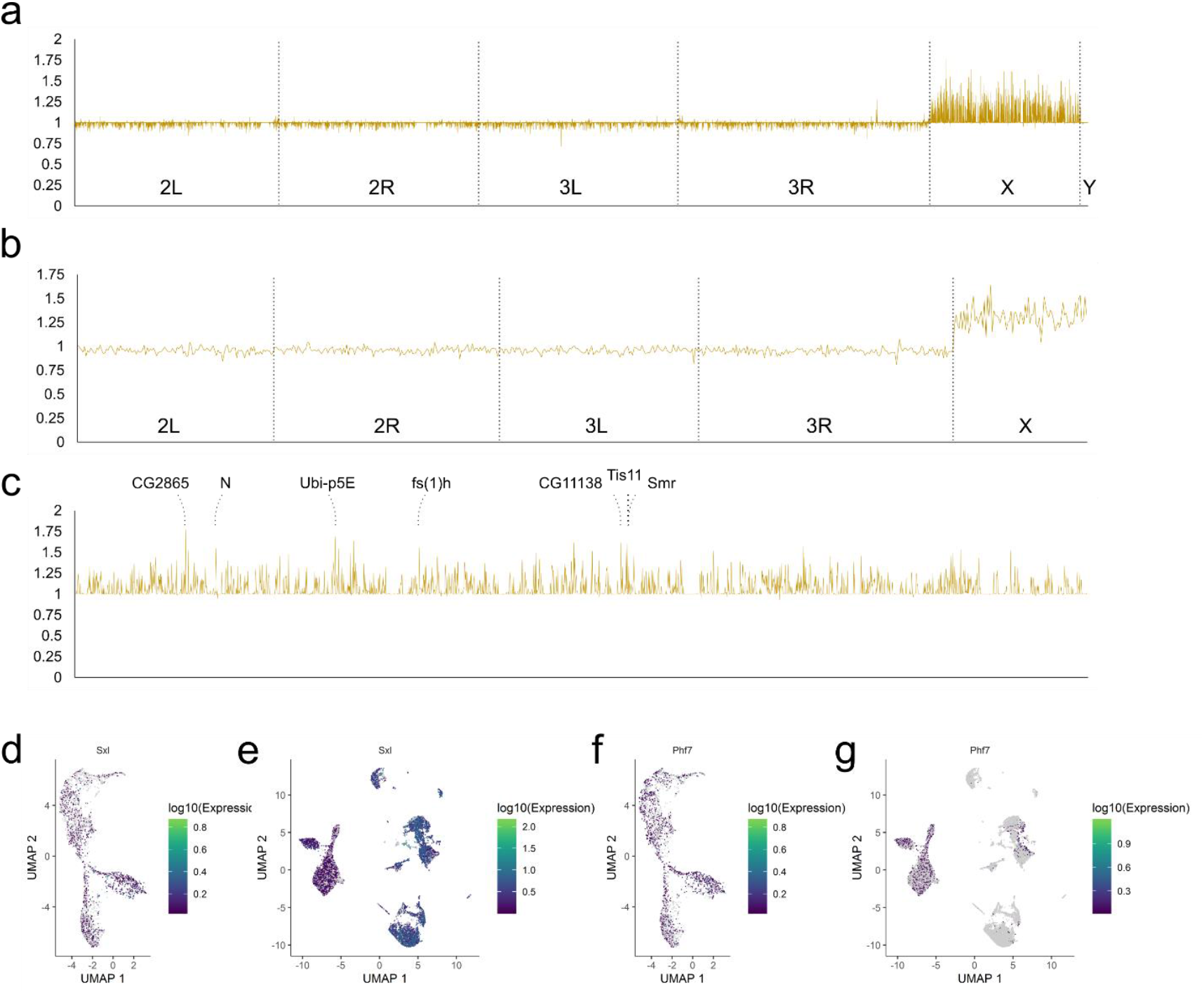
Expression differences between the female and male germline. **a**, Female-to-male expression ratios for genes across all chromosomes from the unsexed dataset. The axes in **a-c** are the same: the X-axis plots the positions on individual chromosomes as indicated and the Y-axis plots the fold differences of expression of female germ cells over male germ cells. **b**, Expression differences between the two sexes for the 647 zygotically activated germline genes from the unsexed dataset. **c**, Sex expression differences from the unsexed dataset highlighted for the X-chromosome. The identities of the highest peaks are indicated. **d-e**, Expression profiles for *Sxl* in the unsexed (**d**) and sexed (**e**) datasets. **f-g**, Expression profiles for *Phf7* in the unsexed (**f**) and sexed (**g**) datasets.

### Candidate genes that control germline sex determination

Our single-cell transcriptome profiles have captured the earliest stages of germline sexual differentiation, and this data enabled us to look for candidate genes of germline sex determination on the X chromosome. From the germline sex expression ratio plots, one can identify individual peaks that are particularly high, and they correspond to greater female-to-male expression ratios (Fig. 4g, Fig. S5c). Interestingly, the identities of these higher peaks between the unsexed and sexed samples match each other very well (Table 1).

**Table 1.** Sex-biased genes in the early germline.

| Top 15 F>M genes<sup>a</sup> |  | Top 15 female marker genes<sup>b</sup> |  | Male marker genes<sup>c</sup> |  |
| --- | --- | --- | --- | --- | --- |
| Unsexed sample | Sexed sample | Unsexed sample | Sexed sample | Unsexed sample | Sexed sample |
| *CG2865*<sup>d</sup> | *MRE16* | *Dsp1*<sup>e</sup> | *fs(1)h* | *vig2* | *vig2* |
| *Ubi-p5E* | *CG2865* | *CG3224* | *Dsp1* | *Hsp26* | *CG2852* |
| *CG3224* | *Ubi-p5E* | *CG11699* | *Ubi-p5E* | *Hsp27* | *stai* |
| *Pits* | *Smr* | *Pits* | *Pits* |  | *p23* |
| *Rpl1215* | *fs(1)h* | *CG2865* | *Smr* |  | *Fer2LCH* |
| *Tis11* | *Pits* | *Rpl1215* | *Imp* |  | *eEF1delta* |
| *Smr* | *N* | *Smr* | *CG11699* |  | *Hsp83* |
| *Dsp1* | *CG32767* | *Tis11* | *N* |  | *mod* |
| *fs(1)h* | *RhoGAP19D* | *CG15891* | *CG32767* |  | *CG1307* |
| *N* | *egh* | *CG14210* | *pico* |  | *wde* |
| *CG15891* | *feo* | *Imp* | *CG15891* |  | *Sod1* |
| *CG14210* | *elav* | *Ubi-p5E* | *arm* |  | *CG15019* |
| *mRpl14* | *Pat1* | *rudhira* | *mRpl14* |  | *IscU* |
| *l(1)G0004* | *Tis11* | *N* | *rudhira* |  | *SeIT* |
| *CG43088* | *Chrac-16* | *fs(1)h* | *Smox* |  | *Nop56* |
|  |  |  |  |  | *MED26* |
|  |  |  |  |  | *ics* |
|  |  |  |  |  | *Prosbeta3* |
|  |  |  |  |  | *Hsp27* |

<sup>a</sup>this category is determined by calculating the female-to-male ratio of normalized expression in cells past the bifurcation point and is ordered by the ratios from high to low
<sup>b</sup>this category lists the top 15 marker genes determined for the female branch ordered from high to low based on their significance
<sup>c</sup>this category lists all male markers with a false discovery rate of <0.05 and is ordered by significance from high to low
<sup>d</sup>the genes highlighted by boxes are common between the unsexed and sexed samples
<sup>e</sup>the genes highlighted in grey encode proteins with potential roles in gene expression regulation

This suggests that these genes are regulated differently from most other genes on the X chromosomes and their higher levels of expression are likely to be functionally significant for sex-specific germline development. Interestingly, there are also a few higher peaks in the sex expression ratio plot of germline prior to bifurcation (Fig. 4c). Among those, two are X-encoded transcription factors Cut and SisA, the latter being a known X-dosage sensor in the soma and recently shown to regulate germline expression of *Sxl*, a known regulator of female germline sexual development (Cline and Meyer 1996; Goyal et al. 2020). These genes may also contribute to germline sex determination.

Another method for identifying sex markers is to call top marker genes for the female and male germ cells past the bifurcation point. We conducted this analysis without taking into consideration transcriptomes of earlier germ cells or somatic cells as germline sex-determining factors would not necessarily have to be germline-or stage-specific. For female germline markers, we focused on the few with the most stringent significance values as many others will show female enriched expression due to incomplete dosage compensation. Of the top 15 female markers, 8 are common between the unsexed and sexed samples (Table 1). Of those 8, 5 also overlap with the genes that exhibit the highest female-to-male expression ratios. Quite intriguingly, several of those such as *fs(1)h, Smrter (Smr)*, and *Pits* encode transcriptional regulators or contain nucleic acid binding domains (Table S4). These genes are prime candidates as sex-determining factors.

**Table S4.**
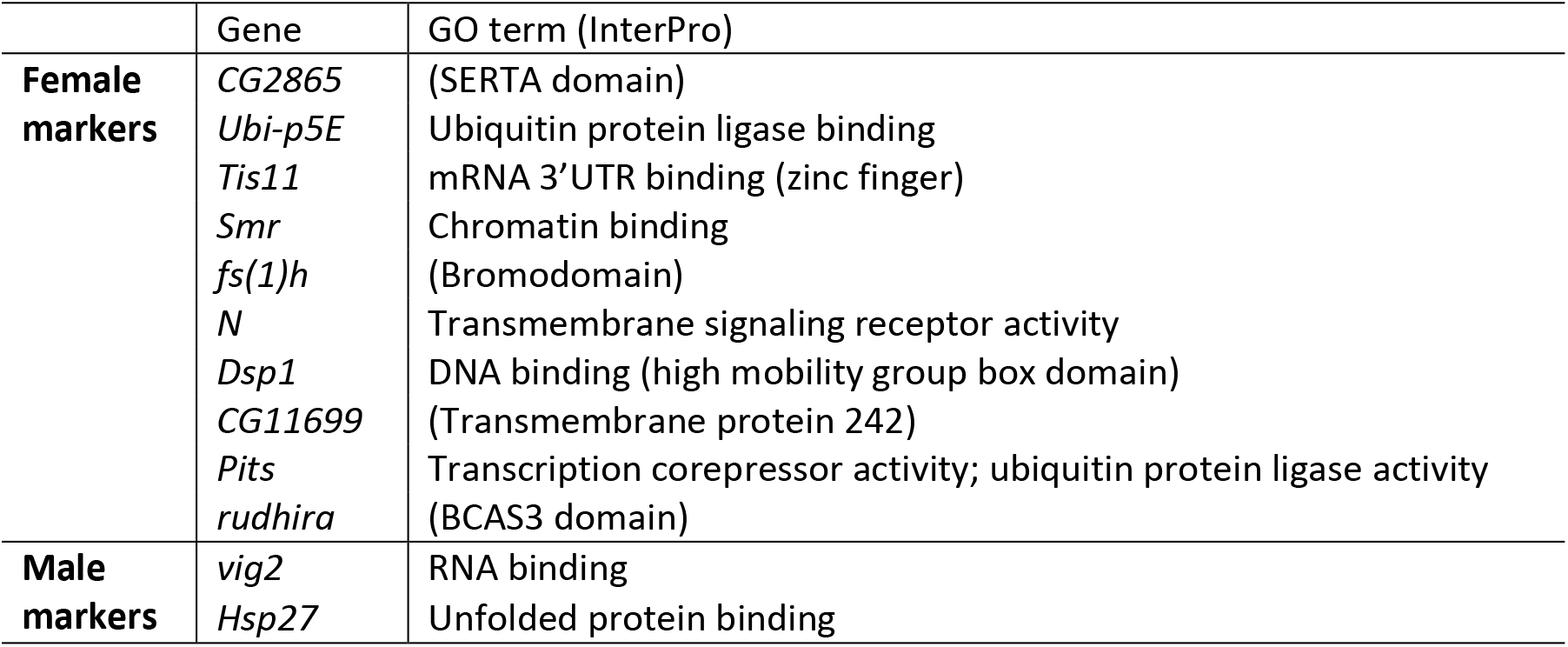
Molecular functions of genes enriched in female or male early germline.

We also looked for male germline markers and found that not only is this list much shorter than that for female markers, the extents of increased expression compared to female germ cells are also smaller (Table 1, Fig. 5a-d). This is possibly due to 8-h male germline being at the cusp of differentiation and thus showing limited male-specific gene activation.

**Figure 5.**
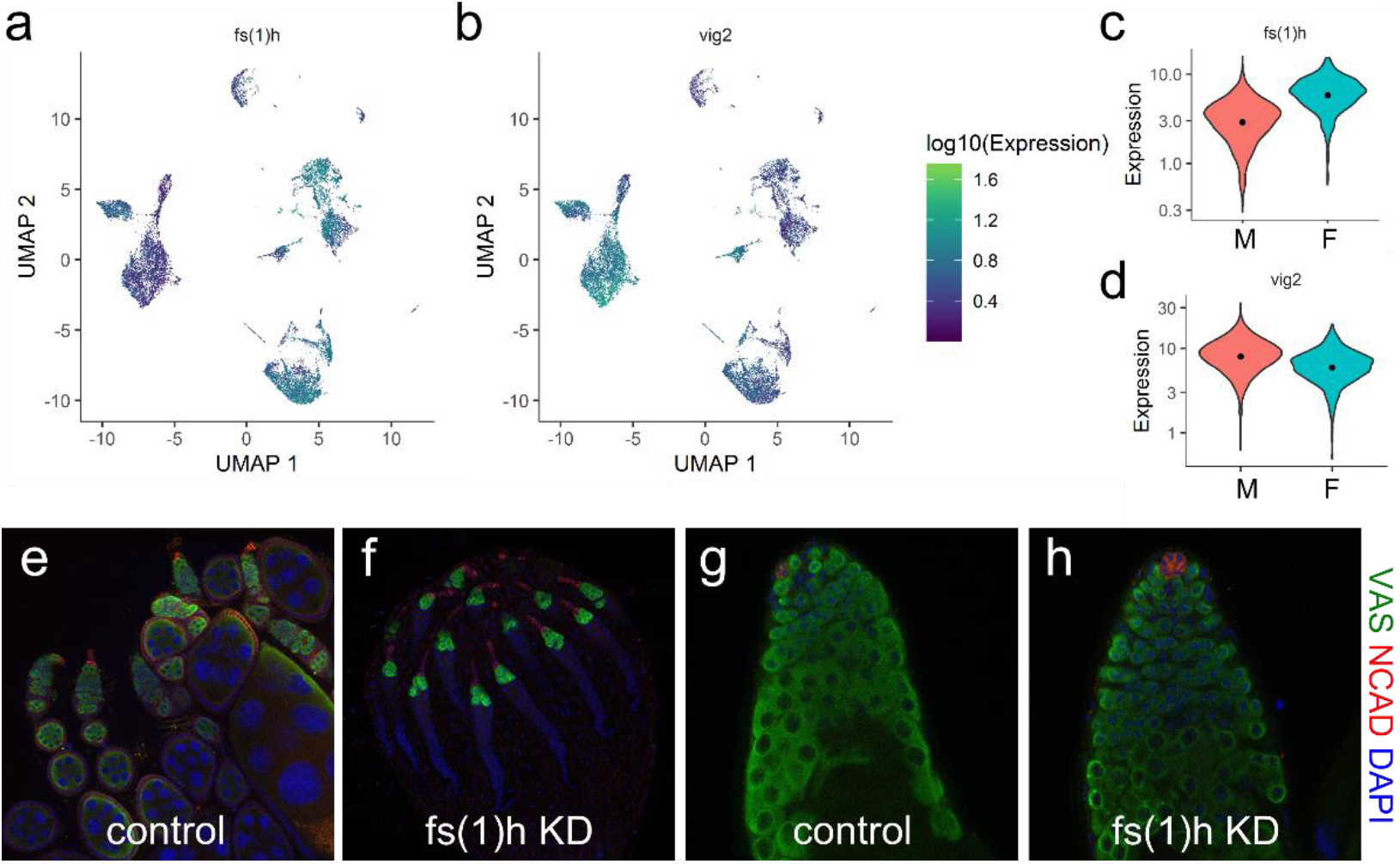
Candidate sex markers in the early germline. **a-b**, Expression profiles of *fs(1)h* (**a**) and *vig2* (**b**) in the sexed dataset. Color codes of the expression levels are indicated to the right. **c-d**, Violin plots of the female marker *fs(1)h* (**c**) and male marker *vig2* (**d**). Orange and turquoise populations indicate male and female expression levels, respectively. **e-h**, Phenotype of RNAi knockdown of *fs(1)h* in the germline of ovary and testis. Samples are stained with antibodies for Vasa (germ cells, green), Cadherin-N (terminal filaments, red), and DAPI (nuclei, blue). **e**, control ovary (*nos-Gal4/+*); **f**, germline knockdown of *fs(1)h* in the ovary (*HSM02723/+;nos-Gal4/+*); **g**, control testis (*nos-Gal4/+*); **h**, germline knockdown of *fs(1)h* in the testis (*HSM02723/+;nos-Gal4/+*).

### A top female marker, *fs(1)h*, regulates female-specific germline development

To investigate whether candidate genes exhibiting sex-enriched expression are important for germline sexual development, we performed functional tests for one of the genes that has consistently ranked highly in our tests of female marker genes, *fs(1)h*. This gene is located on the X chromosome and encodes a bromodomain protein known to cause female sterility. Many studies have demonstrated its functions in gene expression regulation in various contexts (Chang et al., 2007; Digan et al., 1986;

Florence and Faller, 2008), but its possible roles in the germline have not been reported. We tested the functions of *fs(1)h* in female-specific germline development by germline-specific RNAi knock-down (*nos-Gal4, UAS-fs(1)h RNAi*). Immunofluorescence staining of adult testes and ovaries indicate that knocking-down *fs(1)h* expression in the germ cells results in female-specific germline loss (Fig. 5e-h). These results indicate that *fs(1)h* is specifically needed for female germline development. Furthermore, this functional validation indicates that the higher expression of female marker genes are physiologically important and are likely established with unique mechanisms.

## Discussion

Our single-cell transcriptome profiling of the embryonic *Drosophila* germline provides a comprehensive coverage of the molecular signatures at this critical developmental stage. It demonstrates the power of using single-cell techniques to build trajectories that can dissect and reveal characteristics of important lineages that exist in small cell numbers.

### Regulation of germline zygotic activation

Our analyses of TF binding sites suggest that germline-specific gene expression at the time of zygotic activation is regulated largely after transcriptional activation. To investigate these additional mechanisms, we referenced our results with those from a nascent RNA-sequencing study of somatic cells from 3-3.5 h embryos (Saunders et al. 2013). Cross-analyses clearly indicate that polymerase pausing and transcriptional read-through occurs on a substantial number of germline genes in the soma, suggesting that regulation of transcription elongation and RNA stability are important for establishing germline-restricted gene expression. The time window profiled in the Saunders study is a few hours earlier than the somatic cells we analyzed (5-8 hours), but assuming that expression profiles do not change abruptly in the soma within these time frames, we can obtain insightful comparisons between the GRO-seq dataset and our own. In fact, our analyses here may yet underestimate the contribution of polymerase pausing and differential RNA stability to gene expression differences between the soma and germline as the Saunders study was a profile on the entire soma rather the subset that most resembles germ cells in their usage of transcription factors. Future studies that directly compare promoter-proximal polymerase pausing and RNA stability between germ cells and specific somatic cells will clarify the extent these processes are involved in establishing the distinction between the germline and somatic programs.

Based on our scRNA-seq data, transcripts of several TFs and DNA-binding proteins are enriched in the early germline. For those with known binding sites including Caudal (Cad), Dorsal (Dl), and Tailless (Tll), they were not identified in our motif searches, though the possibility of them and the other TFs being involved in zygotic germline activation remains. For genes that encode RNA-binding proteins, there are three that are germline-specific and expressed prior to zygotic activation: *nos*, *oskar*, and *bru1*. Notably, they all have known functions in regulating early germline development (Asaoka et al., 2019; Kim-Ha et al., 1995; Lehmann and Nüsslein-Volhard, 1986). Moreover, there is an example of *nos* regulating stability of a transcript in the embryonic germline (Sugimori et al. 2018). It is quite possible that their roles in germline biology are more extensive than we currently appreciate.

### Sex determination in the germline

The intrinsic information of sexual identity in the *Drosophila* germline comes from its sex chromosome composition via yet unidentified components, and this information would also regulate the process of dosage compensation which does exist in the embryonic germline. Previous studies have reported two germline genes that can induce sex reversal: *Sex lethal (Sxl)* whose expression begins in stage 9 female germline and *PHD Finger Protein 7 (Phf7)* which is expressed starting in stage 13 male germline and on (Hashiyama et al. 2011; Yang et al. 2012). In our data, we observe low expression for *Sxl* and even lower levels for *Phf7* (Fig. S5d-g). Neither are on the top marker lists for the female and male germline. For the female germline, multiple genes are consistently found as top female markers, and several of which are known to or predicted to have functions related to gene expression regulation. These are promising candidates as readers of the germline sex and will be investigated in further functional studies. As the sex expression ratios of these candidates are close to 2-fold, these gene loci most likely harbor characteristics that allow them to escape the germline dosage compensation machinery in the female germline, another very interesting topic for future studies.

Curiously, some genes also exhibit higher female-to-male expression ratios in the germline population prior to the bifurcation point of the sexed. However, to explain how their mRNA levels would become sex-dependent prior to zygotic transcription, one would need to invoke special scenarios such as somatic induction, leaky transcriptional repression in this early phase, or paternal contribution. We will caution that this analysis result stems from a single dataset (the sexed sample) and that the sex expression differences are modest, thus any further investigation will have to start with the validation of sex-dependent expression levels.

In the developmental trajectories, the male branches appear to extend further along pseudotime than the female branches (Fig. 1e, i). This suggests some degree of development present in the male germline at 8-h of embryogenesis and is in line with our knowledge that male germline development precedes that of their female counterparts. Nonetheless, none of our various attempts to find genes that are male-enriched past the bifurcation point resulted in candidates with sufficient confidence beyond *vig2* and heat-shock protein genes as described earlier. We hypothesize that the male germline at this time point is at the very start of development with *vig2* involved in modifying chromatin and heat-shock chaperones being produced to assist folding of the upcoming wave of new proteins.

Our scRNA-seq results portray the early sex-developmental sequence of *D. melanogaster* germline as such: when zygotic transcription is de-repressed, higher expression of select X chromosome genes due to a double dose of X chromosomes drives the germline towards the female program, away from the default male fate. Male germ cells which exhibit lower expression of these female-determining genes initiate development. In contrast, establishment of the female fate would result in the mitotic and developmental quiescence of the germ cells until later larval stages. This is a new model that paves the way for future functional studies on the central germline sex-determining factors.

## Methods

### Fly strains

Fly strains used in this study include *vas-GFP* (Shigenobu et al. 2006) and *Sxl-Pe-EGFP.G G78b* (abbreviated as *Sxl-GFP*, Bloomington Stock Center)*. Sxl-GFP* was used for sexing of embryos. In the first round of scRNA-seq, we used flies that contained four copies of *vas-GFP*. The second round of sequencing was carried out with flies homozygous for the *Sxl-GFP* and *vas-GFP* transgenes, both of which are on the second chromosome.

To perform RNAi knockdown of *fs(1)h* in germ cells, flies carrying the *P{TRiP.HMS02723}attP40* RNAi construct (Bloomington Stock Center) and *nos-Gal4* (Van Doren et al. 1998) transgenes were analyzed.

### Isolation of germ cells

Embryos of the desired age and genotype were collected on grape juice plates, dechorionated, and homogenized with the loose pestle in a Dounce homogenizer for 6-7 strokes. We found that this step alone could release sufficient single germ cells, thus we chose to forgo further enzymatic treatments. The lysates were filtered twice through 70 μm mesh, centrifuged at 850 g for 2 min, and FACS-sorted (FACSAria, BD) to obtain GFP+ germ cells. A small fraction of the sorted cells were examined on a fluorescence microscope (AxioSkop, Zeiss) to document the integrity, purity, and cell number of the resulting samples before being used for library construction on the 10X Genomics single-cell RNA-seq platform and high-throughput sequencing on the NovaSeq 6000 System (Illumina).

We performed two rounds of scRNA-seq experiments. In the first round, we collected 0-4 h and 4-8 h *vas-GFP* germ cells separately as the respective yields for germ cells were quite different (Fig. S1a-d). The purified cells were then mixed in equal numbers and processed via the 10X Genomics drop-seq pipeline (Single Cell 3’ v2). In the second round, we performed scRNA-seq (Single Cell 3’ v3) on sexed 5-8 h germ cells, obtained by FACS-sorting GFP+ cells from 5-8 h *Sxl-GFP, vas-GFP* embryos. 5-8 h female embryos show clear *Sxl-GFP* signals whereas male ones do not, and we hand-separated male and female embryos under fluorescent stereoscopes (Fig. S1e). We validated our sex assignment by examining the sexes of hundreds of sorted embryos after their development to adulthood (data not shown). Embryos that have been separated by sex were subsequently homogenized as described above and sorted by FACS. Compared to the first round (Fig. S1a, c), gating for live cells was further restricted to a subset enriched for germ cells (Fig. S1f, h) to reduce inclusion of somatic cells that may also express GFP from the *Sxl-GFP* transgene (Fig. S1g, i). The purity of the first unsexed sample was close to 100% germ cells as estimated by examining a fraction of the sorted cells by fluorescence microscopy; the purities of germ cells in the sexed female and male samples were estimated to be about 20% and 50% (data not shown).

### Analysis of scRNA-seq data

For the unsexed sample, we obtained sequencing results for 3,810 cells that passed through quality control with the CellRanger software (version 3.0.1, 10X Genomics) with the mean reads per cells being 33,487 and the median genes detected per cell being 3,166. For the sexed samples, 11,001 and 7,222 cells from the female and male samples passed through quality control. The mean reads per cell was 22,241 (female) and 29,231 (male), and the median genes detected per cell was 2,045 (female) and 3,482 (male). 84.9%, 80.1%, and 83.9% of all reads from the three samples, unsexed, female, and male, mapped to the *D. melanogaster* transcriptome (BDGP6.28).

Sequencing results were analyzed with the Monocle3 package to determine clusters and construct a pseudotime that signifies the developmental trajectory of early germ cells (Cao et al. 2019). The data underwent pre-processing (number of dimensions set to 100), dimension reduction using the UMAP method, and clustering with the Leiden algorithm. This gave rise to Y-shaped clusters for germ cells which is determined by expression of two germline markers, *nos* and *vas*. To order cells in the germline clusters, we chose the “root” to be at the end of the stem of the Y-shaped clusters based on the expression patterns of *pgc* and *gcl* (Fig. S2c, d) which consequently enables assignment of pseudotime in the germline clusters.

To investigate expression trends, we determined expression modules using default parameters except for the resolution being set to 0.0001. To identify markers of designated clusters, we utilized the top_markers function to find top genes that delineate various cell populations and used *q* values or marker scores in addition to prioritize candidate genes and adjust stringency. For the determination of germline marker genes, a 0.3 marker score cut-off was used. We noticed a few genes in the marker lists that overlap, especially when comparing the maternal and zygotic germline marker lists. When we examined the expression profiles of duplicated entries, most of them had relatively high expression throughout pseudotime. These genes were a minority on the lists and were not included in the list of zygotically activated genes. To generate the subsets of germline markers whose expression is highly germline-enriched, we eliminated genes whose average somatic expression was greater than 0.03 as determined by calculating the average expression of all somatic cells included in the sexed sample.

To examine expression progression of specific genes, we selected the germ cells in the stem of the Y-shaped clusters as well as in the male branch as these cells together represent a linear developmental progression along pseudotime. This allowed us to graph changes in expression of individual genes as a function of pseudotime.

### Analyses of marker gene characteristics

To identify pathways that are enriched in the zygotically activated germline genes, we performed ordered queries for KEGG Pathway and GO Term: Cellular Components analyses using the g:Profiler platform (biit.cs.ut.ee/gprofiler/gost, Raudvere et al., 2019) with false discovery rate thresholds of 0.05.

To look for enrichment of transcription factor binding sites in the zygotic germline genes, we used the g:Profiler interface to search the 1 kb region upstream and downstream of the transcription start sites (TSSs) of candidates for binding sites catalogued in the TRANSFAC database (Release 2019.3, classes: v2) with *p* < 0.05. For *de novo* identification of sequence motifs enriched in the promoter regions of zygotic germline genes, we extracted the 1 kb region upstream of TSSs of all soma-positive and negative zygotic germline genes from the Ensembl BioMart database and used the MEME-suite for motif discovery using the 0-order model for background correction and with the statistical significance (E-value) cutoff of 0.0001 (Bailey et al. 2009). To examine the distribution of motifs in markers of different clusters, we used the FIMO tool for motif scanning of match sites in the 1kb region upstream of TSSs of germline genes or top 100 markers of clusters 2-4 with *p* < 0.0001 (Grant et al. 2011).

To examine the extent of polymerase pausing in the embryonic soma, we referenced pausing indices from the Saunders study that performed GRO-seq for 3-3.5 h embryos (GSE41611, Saunders et al., 2013). To determine transcript stability based on the Saunders’ study, we referenced a re-analysis which was mapped to the newer *Drosophila* genome assembly (dm6, GSM3281693, GSM3281694) to calculate mapped reads per bin (MRPB) in the gene body regions defined as being from +100 to the end of each gene. This is to avoid reads that reflect promoter-proximal polymerase pausing rather than transcriptional read-through. MRPB values for gene body regions were determined by subtracting the MRPB values in the first 100 bps downstream of TSSs from those for the entire length of genes, all computed with multiBigwigSummary (version 3.3.2.0.0) from the deepTools2 package via the Galaxy platform and corrected for their relative lengths (Goecks et al. 2010; Ramírez et al. 2014). The lowest MRPB values were used in subsequent analyses for genes with more than one isoforms. The gene body MRPB values were further divided by the average steady-state expression levels calculated for all somatic cells combined based on our own scRNA-seq data to obtain indices that reflect RNA stability. The Wilcoxon rank sum test was used to determine statistical significance comparing RNA stability of zygotic germline soma-negative and positive genes. The Kruskal-Wallis test followed by Dunn’s multiple comparison test was used to compare RNA stability of soma-negative and positive germline genes that exhibit polymerase pausing and not.

To calculate the gene-by-gene female-to-male expression ratios, we used the average expression of each gene in all germ cells in the male sample, all germ cells in the female sample, all somatic cells in the male sample, and all somatic cells in the female sample. Counts normalized by log-transformation were used as expression values for each gene.

### *In situ* hybridization and immunofluorescence staining

*In situ* hybridization chain reactions (HCR v3.0, Molecular Instruments) were performed as recommended by the vendor on 0-16 hr (25°C) embryos to validate scRNA-seq results. Embryos were dechorionated with 100% bleach for 2 min prior to fixation in 4.5% formaldehyde and clearing with xylene substitute (Sigma-Aldrich) to minimize auto-fluorescence. Subsequently, samples were hybridized with 2 pmol of split-initiator probes overnight at 37°C to detect mRNA targets, then incubated with 6 pmol of hairpins labeled with various fluorophores overnight at room temperature to generate fluorescent amplification polymers. The embryos were then stained with a rabbit-α-Vasa antibody (d-260, 1:250, Santa Cruz Biotechnology) followed by an Alexa Flour 488-conjugated goat-α-rabbit secondary antibody (1:1000, Jackson ImmunoResearch) to mark the germ cells. Confocal images were taken on a LSM780 (Zeiss).

Immunofluorescence was performed by fixing gonads from 1-3 day old adult flies for 15 min in 4% formaldehyde and overnight staining of antibodies at 4°C. Primary antibodies used were rabbit-α-Vasa, rat-α-Cadherin-N (1:20, EX-8, DSHB); secondary antibodies used were conjugated with Alexa Flours. Samples were also stained with DAPI (1 μg/ml) to mark nuclei before imaging on Apotome.2 (Zeiss).

## Data Access

The scRNA-seq datasets described in this study are deposited at the NCBI Gene Expression Omnibus (GSE150568).

## Acknowledgements

The authors would like to thank the Yang lab for technical assistance, and Mark Van Doren and the lab of Brian Oliver for input on the project. Services of scRNA-seq and de-multiplexing of the sequencing reads were provided by BioTools. Flow cytometry and confocal microscopy was performed at the Core Facility and Imaging Center at Chang Gung University. This work was funded by a MOST grant from Taiwan (108-2628-B-182-007) to S.Y.Y. and Chang Gung Memorial Hospital Grants to S.Y.Y. (CMRPD1K0231) and S.D.F. (CMRPD1J0251).

## Author Contributions

Conceptualization, S.Y.Y; Data acquisition, H.C.C., H.W.L, H.H.H., S.Y.Y.; Data analysis, S.Y.Y., Y.R.L.,S.D.F.; Writing, S.Y.Y.,S.D.F., Funding, S.Y.Y., S.D.F.

## Supplementary Tables

Tables S1 and S2: see complete Excel files online.

